# plantiSMASH 2.0: improvements to detection, annotation, and prioritization of plant biosynthetic gene clusters

**DOI:** 10.1101/2025.10.28.683968

**Authors:** Elena Del Pup, Charlotte Owen, Ziqiang Luo, Hannah E. Augustijn, Arjan Draisma, Guy Polturak, Satria A. Kautsar, Anne Osbourn, Justin J.J. van der Hooft, Marnix H. Medema

**Affiliations:** Bioinformatics Group, Wageningen University, Wageningen, the Netherlands; John Innes Centre, Colney, Norwich, UK; Institute of Biology, Leiden University, Leiden, the Netherlands; Institute of Plant Sciences and Genetics in Agriculture, The Hebrew University of Jerusalem, Rehovot, 7610001, Israel; DOE Joint Genome Institute, Lawrence Berkeley National Labs, Berkeley, CA 94720, USA; Department of Biochemistry, University of Johannesburg, Johannesburg 2006, South Africa

**Keywords:** genome mining, transcriptional regulation, plant specialized metabolism, natural product discovery, biosynthetic gene cluster

## Abstract

Plants produce bioactive compounds as part of their specialized metabolism, with applications in medicine, agriculture, and nutrition. The biosynthesis of a growing number of these specialized metabolites has been found to be encoded in biosynthetic gene clusters (BGCs), creating increasing demand for genome mining tools to automate their detection. plantiSMASH enables the identification of putative plant BGCs through a rule-based approach, available via both command-line and web interfaces. Here, we present plantiSMASH 2.0 (https://plantismash.bioinformatics.nl/), a major update that expands and improves the original framework with revised and additional BGC detection rules (now supporting 12 BGC types), substrate prediction for selected enzyme families, and regulatory analysis through transcription factor binding site detection. The updated plantiSMASH 2.0 database includes 30,423 putative BGCs across 430 genomes. Together, these improvements make plantiSMASH 2.0 a powerful and comprehensive platform for the detection and characterization of plant biosynthetic pathways, supporting and accelerating research in plant specialized metabolism and plant natural product discovery.

## Introduction

Plants produce an extraordinary diversity of specialized metabolites that mediate essential ecological functions such as defense against herbivores and pathogens, attraction of pollinators, and allelopathy [1,2]. These metabolites are also very relevant for human use, serving as a source for pharmaceuticals, agrochemicals, and nutraceuticals [3,4]. Unlike primary metabolites, specialized metabolites typically exhibit lineage-specific distribution and structural diversity, shaped by dynamic evolutionary processes including gene duplication, neofunctionalization, and horizontal gene transfer [5,6].

A growing body of work has revealed that the biosynthesis of many specialized metabolites produced by plants is encoded by physically co-localized genes, so-called biosynthetic gene clusters (BGCs). While BGCs have long been recognized and extensively studied as a pervasive feature of microbial genomes, advances in plant genomics and metabolomics have recently demonstrated the presence and importance of BGCs in plant specialized metabolism [7]. Plant BGCs often encode not only core biosynthetic enzymes such as terpene synthases or polyketide synthases but also tailoring enzymes and occasionally transporters and transcriptional regulators, partly mirroring the modularity seen in microbial BGCs.

Genome mining strategies that computationally predict BGCs from genomic data have significantly accelerated the discovery and characterization of these clusters. The antiSMASH platform has emerged as a foundational tool in microbial BGC research, employing rule-based annotation based on biosynthetic enzyme-coding gene identification using profile Hidden Markov Models (pHMMs) [8]. Previously, plantiSMASH 1.0 [9] was developed as an extension of antiSMASH comprising a rule-based detection framework tailored to plant biosynthetic enzymes. plantiSMASH uses manually curated BGC detection rules based on experimentally validated plant BGCs that define which core biosynthetic functions need to exist in a genomic locus to constitute a potential (putative) plant BGC. While it is difficult to estimate the percentage of these loci that are bona fide BGCs (previous evidence suggests that at least a good proportion of them are coexpressed [9]), identifying these loci systematically then allows them to be further analyzed based on e.g. coexpression data or integrative omics to prioritize them for experimental characterization. plantiSMASH is available both as standalone software and a web service, which includes precalculated results for 48 species that can be browsed and downloaded. The software accepts annotated genome files (GenBank or FASTA+GFF) to identify BGCs and generates a comprehensive interactive summary to aid prioritization. Since its 1.0 release in 2017, plantiSMASH has become the leading platform for plant genome mining and has been widely used in studies to identify new plant BGCs across diverse plant lineages [10–13]. The CoExpress module in plantiSMASH allows for coexpression analysis between detected BGCs and with other genes across the genome, thereby supporting the identification of partially clustered biosynthetic pathways. With over 50 plant BGCs experimentally characterized across plant species to date [14], including novel BGC families, there is continued interest in optimizing genome mining strategies to automate their detection. Furthermore, both declining sequencing costs and increasing genomic availability continue to increase the potential and applicability of genome mining approaches [15]. This calls for plantiSMASH to include BGC detection rules up to date with current biochemical knowledge.

Here, we present the updated plantiSMASH 2.0 (https://plantismash.bioinformatics.nl/), which significantly expands and improves upon the previous version. BGC detection rules were revised and expanded to incorporate new experimentally validated cluster types characterized since the initial release. Additionally, we expanded the plantiSMASH database with precalculated results from the original 49 genomes in version 1.0 to 430 genomes, thereby reflecting newly sequenced species and updated genome assemblies. Updating the database enhances the accuracy and coverage of similarity-based analyses performed via clusterBLAST. Furthermore, we included prioritization of putative BGCs via functional and regulatory evidence, implementing substrate prediction and transcription factor binding site (TFBS) detection. Substrate specificity prediction has been introduced for a set of characterized enzyme subfamilies, expanding functional annotation of BGCs and associating them with putative products. We implemented TFBS detection to prioritize putative BGCs for experimental validation, facilitating a deeper understanding of their regulatory context. Finally, we updated the software codebase and provided extensive documentation to facilitate co-development.

## Materials and Methods

### Code optimization, documentation, and maintainability

To enhance the performance, maintainability, and usability of plantiSMASH, the entire codebase was migrated from Python 2 to Python 3. Dependencies have been updated for compatibility. To support reproducibility and long-term accessibility, releases of plantiSMASH are now archived and versioned with a persistent DOI via Zenodo (https://doi.org/10.5281/zenodo.15412177). This enables traceable, FAIR dissemination in line with best practices for research software [16].

plantiSMASH is now available as a containerized web application via Docker, improving reliability and scalability of the web service, simplifying deployment for local or institutional hosting, and ensuring consistent runtime environments. Additionally, a comprehensive documentation and installation instructions are now available (https://plantismash.github.io/documentation/), and guides to build the application Docker images are provided on the repository (https://github.com/plantismash/plantismash-docker). These distribution methods reduce setup time, eliminate dependency conflicts, and make the tool easily accessible across platforms. A wiki for developers (https://github.com/plantismash/plantismash/wiki) has been introduced for advanced users to define custom detection rules, subfamily prediction modules, and clusterBLAST databases.

### New BGC types for an extended range of plant species

We collected 87 experimentally characterized BGCs from the literature and selected 28 to update the plantiSMASH detection rules that capture them (Table S1), including the 40 characterized BGCs available in MIBiG 4.0 [17]. In plantiSMASH 2.0, the number of supported BGC types increased from six to twelve, including new detection rules for fatty acid, transporter-associated, sesterterpene, strictosidine-like, phenolamide, and cyclopeptide BGCs (Table S2). An additional 43 domain profiles are introduced for BGC detection, including 15 additional non-core (generic) domains (Table S3). Cyclopeptide BGCs are identified through the co-occurrence of BURP domains and repeat motifs, as detailed in a previous study [18]. Motif detection for BURP-associated domain repeats is displayed in the plantiSMASH output, as shown in Figure 1B. To support repeat detection in the case of ambiguous amino acids, the cyclopeptide detection module incorporates an averaging strategy across uncertain amino acid positions. New profile Hidden Markov Models (pHMMs) have been added to facilitate specific detection of signature genes for other BGC types, such as those for ornithine and tyrosine decarboxylases, from the detection rule of phenolamide clusters in rice [19,20]. In addition to the expanded rule set, we increased the number of genomes with precomputed plantiSMASH results from 49 in version 1.0 to 430 in version 2.0 (Table S4). We assessed 440 available annotated reference Streptophyta genomes hosted by NCBI (https://www.ncbi.nlm.nih.gov/datasets/genome/), removing those with poor annotation or assembly quality (<10,000 protein-coding genes, or contig-level assemblies with <10 BGCs in plantiSMASH 1.0). An additional nine genomes from the Gentianales and Poeae were included for coverage of key known BGCs.

**Figure 1.**
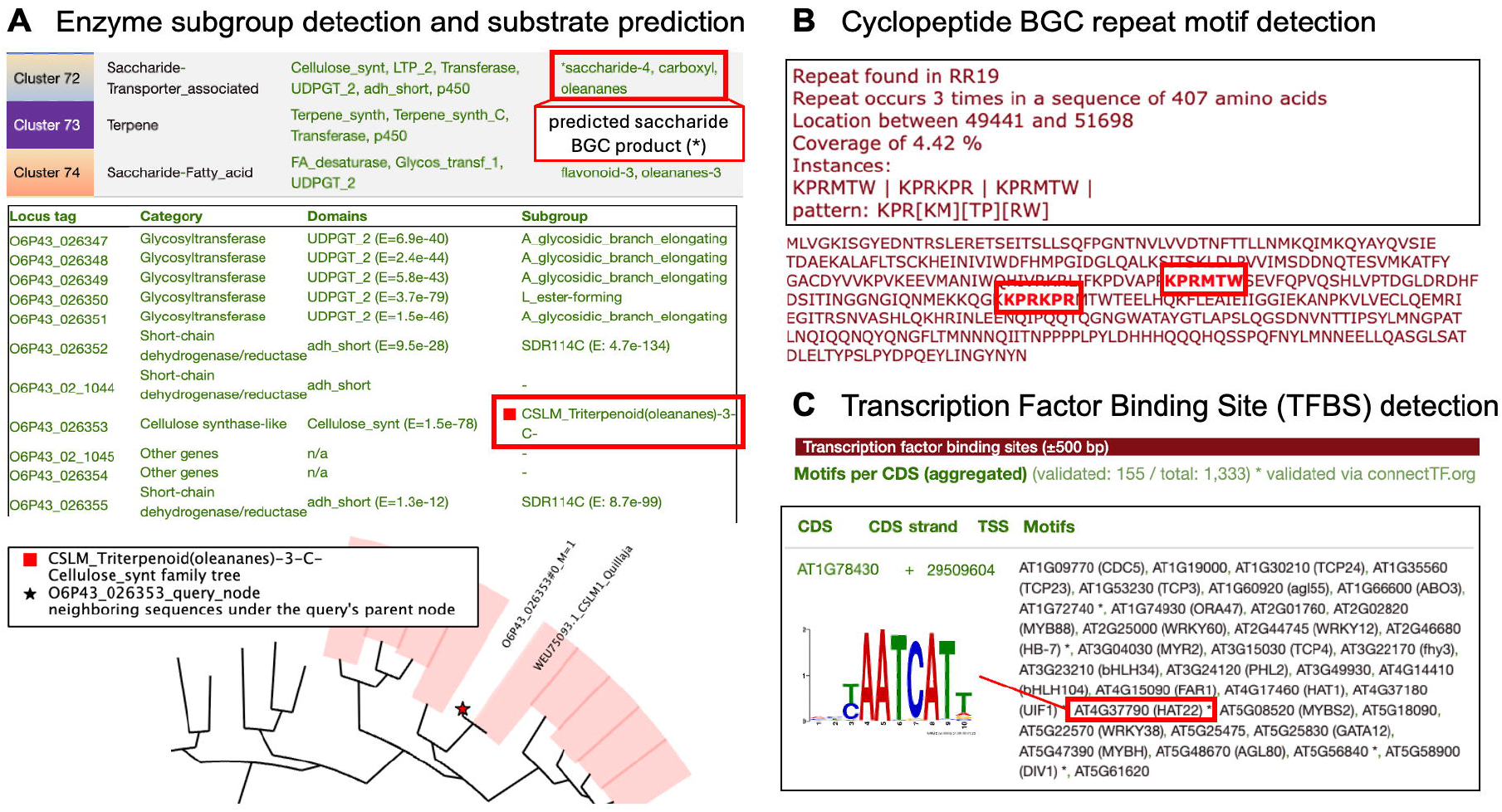
New features and modules in plantiSMASH 2.0. A. Enzyme subgroup detection module and BGC substrate prediction, domain subgroups, and phylogenetic placement in BGC72 in *Quillaja saponaria*, corresponding to the QS-21 locus. B. Detection of cyclopeptide BGCs via repeat motif detection. C. Detection of Transcription Factor Binding Sites (TFBSs) and related output in *A. thaliana*.

### Enzyme subfamily assignment and BGC substrate prediction

To facilitate the prediction of putative products of detected BGCs, we included a module that detects enzyme substrate classes associated with enzyme subfamilies. Currently supported enzyme subfamily predictions are based on curated literature for cellulose synthases (CSLs) [21,22], UDP-glucuronosyltransferases (UGTs) [23], short-chain dehydrogenases/reductases (SDRs) [24], and oxidosqualene cyclases (OSCs) [25]. Each enzyme family’s detection rules and associated HMM profiles have been integrated into plantiSMASH to recognize subfamily-specific signatures. Enzymes are assigned to subfamilies based on their sequence similarity via subgroup-specific HMM profiles [26] and evolutionary relationships inferred via the phylogenetic placement tool *pplacer* for CSLs, UGTs, and OSCs [27]. An E-value for the HMM-detected subgroup is shown when there is disagreement with the subgroup assignment via *pplacer*. Phylogenetic assignment shows the subgroup placement of the query enzyme and then displays the name of the closest leaves to the query, to support further functional exploration (**Figure 1A**). In the BGC overview, ‘*’ marks product types consistent with the core enzyme functions. The substrate prediction module supports user-defined custom subgroups for additional enzyme families based on custom phylogenetic inference and substrate characterization.

### Transcription factor binding site prediction

To enhance the functional annotation of regulatory elements within putative BGCs, we implemented a transcription factor binding site (TFBS) prediction module, analogous to the TFBS Finder module introduced in antiSMASH 7.0 [8]. Binding site identification leverages validated plant TFBS motifs from the PlantTFDB database [28]. The quality of identified motifs was ensured through over-dominance control (≤0.55 average per-position mass of any single nucleotide), and viability (p-value=1×10^−5^). Additionally, we manually curated motifs based on visual inspection of information content and structure, such as the presence of palindromes, according to the method illustrated in LogoMotif [29] (See Supplementary Fig. S1 and Data availability). The TFBS Finder module uses motif position weight matrices (PWMs) to annotate putative TFBSs in a user-defined window upstream of the gene transcription start site and 50 bp downstream, via the MOODs scanner [30]. Additionally, hits are mapped to validated interactions between 662 TFs and target genes in ConnectTF [31] (indicated with ‘*’ in the output summary) (Figure 1C). Common names for the TFs are obtained from TAIR [32]. TFBS Finder results for *Arabidopsis thaliana* are available in the plantiSMASH 2.0 database, with 1*10^-4^ p-value and 500 bp window scanning size (See Data availability). Enrichment analysis of TFBS families across BGC types is performed in *A. thaliana*.

## Results

### Advancing the detection of new BGC types and their products

Across 430 species in the plantiSMASH 2.0 database, 30,423 putative BGCs were detected using the updated rules set, compared to 17,856 putative BGCs in plantiSMASH 1.0 (**Figure 2**). These are available on the plantiSMASH database (https://plantismash.bioinformatics.nl/precalc/v2/). Similarity to all 40 experimentally characterized plant BGCs from MIBiG 4.0, together with the avenacin and QS-21 loci, is detected via the knownclusterBLAST module. The clusterBLAST database was expanded from 7,978 to 135,976 query BGCs, leading to the identification of 101,336 similar subject clusters. Cyclopeptide BGC detection included BGCs with repeat motifs occurring outside the BURP enzyme, resulting in 14.33% (4,361) putative cyclopeptide BGCs in the plantiSMASH 2.0 database. Comparison of cyclopeptides BGC counts with internal and external repeats to the BURP enzyme is noted in Figure 2.

**Figure 2.**
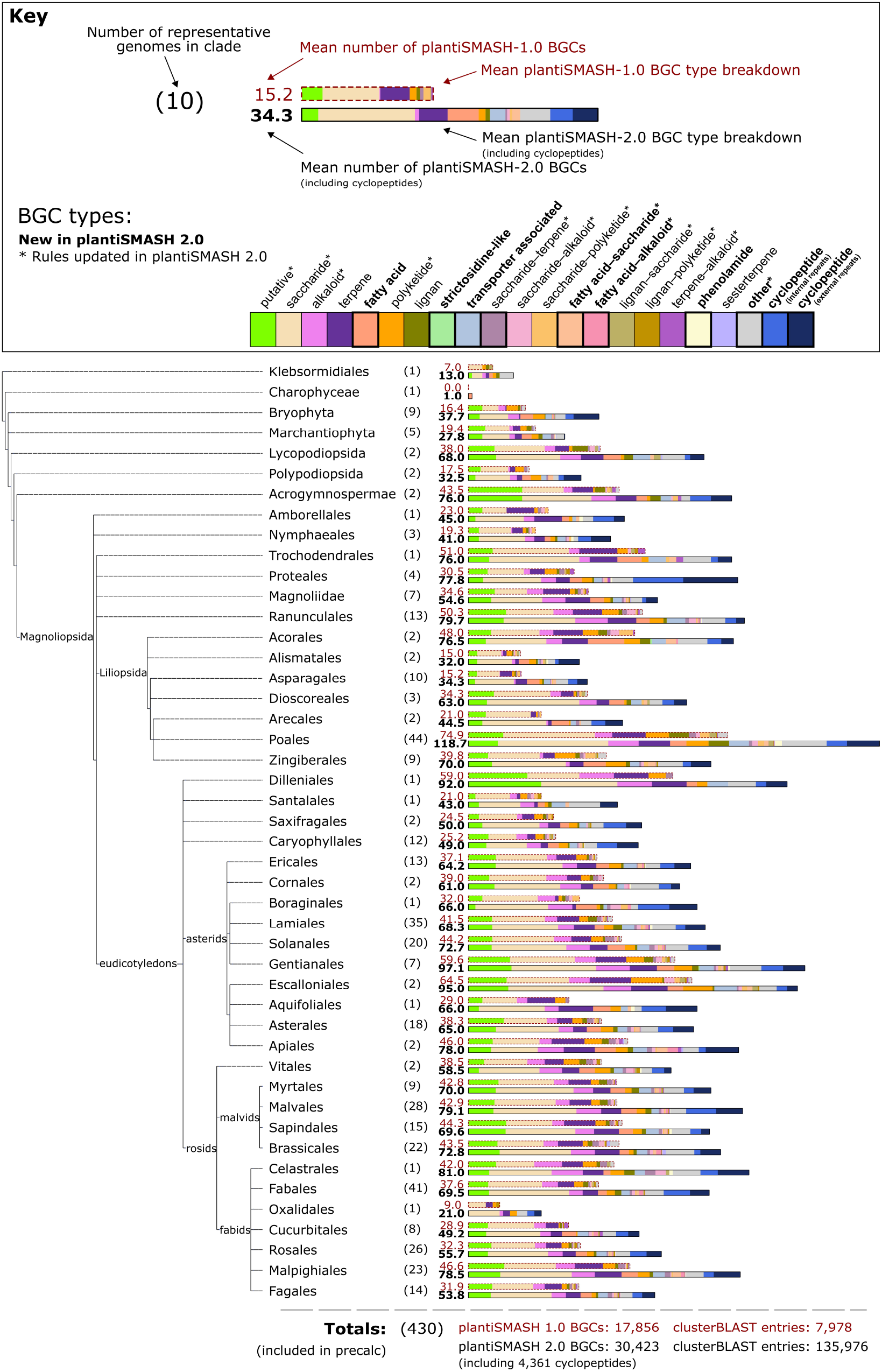
Summary of BGCs detected in plantiSMASH 2.0 compared with plantiSMASH 1.0 across 430 Streptophyta genomes grouped by clade. A detailed overview of putative BGCs per species is available online on the plantiSMASH database 2.0 (See Data availability).

With updated rules, BGCs that have been characterized since the publication of plantiSMASH 1.0 are now identified, such as the QS-21 locus in *Quillaja saponaria* [10,33] and the strictosidine locus in *Catharanthus roseus* [34] (Figure 3A). Strictosidine-like BGCs were also detected in *Gelsemium sempervirens* (as previously characterized [35]), but also in species such as *Acorus calamus* and *Spinacia oleracea*. In *Q. saponaria*, we detected three other saccharide BGCs with partial similarity to the QS-21 locus, as well as predicted associated saccharide products identified by enzyme subfamily detection, containing genes characterized as part of the saponin biosynthetic machinery [10].

**Figure 3.**
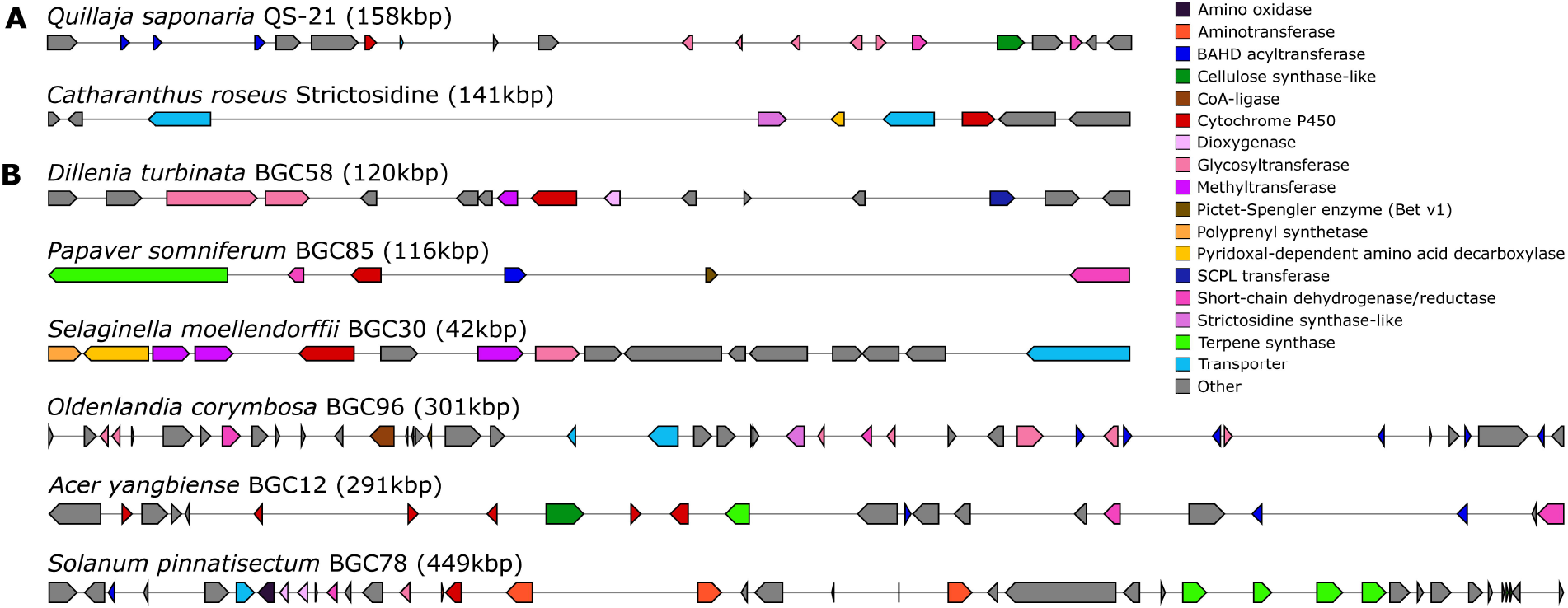
Examples of BGCs detected by plantiSMASH 2.0. A. Examples of recently characterized BGCs now detected in plantiSMASH. B. Examples of uncharacterized detected candidate BGCs.

The new clusterBLAST database and knownclusterBLAST against MIBiG 4.0 are a powerful way of rapidly assessing BGC similarity and potential function. For example, the core of the experimentally characterized avenacin cluster in *Avena strigosa* consists of 12 genes [36]. The plantiSMASH 2.0 analysis detected a BGC corresponding to the avenacin cluster locus, much larger than the core BGC (CD-HIT=27), as it also satisfies the detection rules for other BGC types. In plantiSMASH 1.0, these regions were defined as three separate BGCs at the chromosomal tip. Homologous BGCs identified by clusterBLAST and knownclusterBLAST in *A. strigosa* and related species are shown in Supplementary Fig. S2.

### Improving functional BGC annotation by substrate detection and product prediction

Subgrouping of key gene families allows for improved prediction of products from BGCs. For example, the QS-21 locus is automatically assigned with 1) group A and L UGTs (indicative of glycosidic branch elongation and ester-bond formation), 2) the cellulose synthase-like M subgroup required for the first glucuronidation (specific to oleanane scaffolds), and 3) the SDR114C subfamily required for D-Fucose formation [10]. These automatic assignments, when coupled with the BGC-type and clusterBLAST outputs, allow for quick and powerful assessment of putative clusters of interest.

Figure 3B shows some example uncharacterized clusters provided in the plantiSMASH 2.0 database. BGC12 from *Acer yangbiense* and BGC85 from *Papaver somniferum* both contain OSCs predicted to make beta-amyrin or similar scaffolds, and are clustered with P450s, acyltransferases, and four different subgroups of reductases (SDR110C, SDR7C, SDR460A, and AT4G33360.1-like) - all suggesting potential for production of an acylated triterpene product. Furthermore, the *A. yangbiense* locus also contains a cellulose synthase-like M gene, strongly indicating a glucuronidation at the C-3 position as seen in e.g. QS-21 [10]. ClusterBLAST for these BGCs also shows homologous sequences in *Acer sp*. and *Papaver sp*. The strength of the clusterBLAST module in evaluating likely functional conservation of BGCs between species is especially apparent when multiple reference genomes are available across a clade. For example, BGC78 from *Solanum pinnatisectum* not only has the BGC type and subgrouping indicative of a glycoalkaloid and a knownclusterBLAST hit on alpha-solanine/chaconine (MIBiG entry BGC0002722), but also has clear similarity to loci in other *Solanum spp*. (Supplementary Fig. S3).

In cases where there is no clear homolog to a known BGC, the improved annotation helps users assess potential function. For example, BGC96 from *Oldenlandia corymbosa* encodes a strictosidine synthase-like enzyme, BAHD acyltransferases, UGTs (groups O, E, and A), reductases (SDR114C and SDR6E), transporters (MatE and ABC), a Pictet-Spenglerase, and CoA-ligase genes – all potentially involved in the production of a glycosylated, acylated alkaloid. BGC58 from *Dillenia turbinata* appears to be an ‘accessory’ BGC, containing enzymes typically involved in scaffold decoration of a triterpenoid or flavonoid (two group D UGTs, a dioxygenase, a P450, a SCPL acyltransferase, and a methyltransferase). *Selaginella moellendorffii* is an early vascular plant that contains clusters with no similarity to known BGCs, such as BGC30, which encodes O-methyl transferases, amino acid decarboxylases, a polyprenyl synthetase, a group L (i.e., ester-forming) UGT, a P450, and an ABC transporter. As such, it potentially encodes the machinery to produce a prenylated indole alkaloid glycoside.

### Prioritizing BGCs with transcriptional regulation via binding motif detection

From the 619 experimentally validated PWMs from PlantTFDB [28] in *A. thaliana*, 576 PWMs passed the quality filters, and 354 PWMs were kept after manually curating an exclusion list (See Data availability). Across 576 PWMs, we obtained 589 strong hits validated with ChIP-seq for all 65 *A. thaliana* detected BGCs, while across the 354 manually curated motifs, we obtained 405 strong hits validated with ChIP-seq (Supplementary Fig. S4-5). The occurrence of G-box-like motifs (CACGTG) is predicted to result in many overlapping sets of binding sites and is known to bind to many TF families [37]. Enrichment analysis of motifs by TF family across BGC types, for the manually curated motifs, helps find biologically significant regulatory patterns and exclude noise from motif families that are predicted to have very high occurrence across all BGC types. Across *A. thaliana* cluster types, our enrichment analysis of the manually curated PWM set revealed strong hits with ChIP-seq evidence across the NAC, WRKY, MYB, bZIP, and ERF TF families (Supplementary Fig. S6-7). This pattern mirrors the known regulatory logic of specialized metabolism in *A. thaliana*. WRKY33 is known to activate the genes for camalexin production, an indole-derived phytoalexin with a critical role in defense, explaining the signal for the WRKY family in alkaloid/polyketide-alkaloid-type BGCs clusters [38]. APETALA2/ethylene-responsive (ERF/AP2) factors are observed in alkaloid-type BGCs, consistent with their role in frequently regulating defense-linked specialized pathways and cross-talk with jasmonate and ethylene metabolism [39]. bZIP (e.g., ABF/AREB, TGAs) enrichment in specialized metabolism is connected to stress-driven metabolic reprogramming [40]. MYB and NAC regulators that govern the phenylpropanoid network are enriched in lignan-type BGCs [41]. No significant enrichment for the *A. thaliana* BGCs is found for the bHLH family of regulators, which is linked to jasmonate-driven specialized metabolism, driving glucosinolate biosynthesis [42]. Also, the TCP family of TFs is not enriched in current BGC types, despite playing key roles in regulating the biosynthesis of specialized metabolites [43].

## Discussion

plantiSMASH 2.0 greatly expands the capacity for automated plant genome mining by detecting a wider range of putative BGCs, including experimentally characterized ones, and supporting the functional characterization of the uncharted plant chemical space. The integration of both substrate prediction, by enzyme subfamily assignment, and TFBS detection enhances interpretability and downstream analyses of detected BGCs. With over 30,000 mined BGCs in the plantiSMASH 2.0 database, users can search for specific BGC types and gene families, inspect subgroup and clusterBLAST outputs, and comprehensively assess and prioritize genes for study across the whole plant kingdom. Improvements over the newly introduced modules will rely on future studies on enzyme family phylogenetics and subgroupings, as well as the validation of motifs of regulatory elements in other species beyond *A. thaliana*. Specialized metabolism in plants is also regulated by signatures other than transcription factors. For instance, chromatin remodeling factors such as ARP6, histone mark H3K27me3, and histone variant H2A.Z, play a role in the regulation of thalianol and marneral clusters [44]. Future work can focus on including these other features of specialized metabolism regulation.

Detection rules in plantiSMASH are based on manually curated criteria, representing valuable starting points that can be iteratively enhanced through continued experimental validation and community-driven curation efforts. However, not all enzyme families involved in specialized metabolism currently have comprehensive or fully characterized HMM profiles, providing exciting opportunities for further investigation and refinement of annotations. Addressing these challenges through ongoing experimental characterization and expansion of both HMM profiles and rule-based BGC annotations will be crucial to fully capture the diverse biosynthetic capabilities encoded within plant genomes.

While plantiSMASH is a powerful tool to explore specialized biosynthesis encoded in plant genomes, unclustered or partially clustered biosynthetic pathways that frequently occur in plants with their complex genomic arrangements are difficult to capture with genomic detection rules alone. Detection of these pathways must take advantage of complementary approaches for pathway prediction, such as relying on gene coexpression patterns, available in plantiSMASH via the CoExpress module, and associating them ultimately with metabolomics datasets. Gene expression data is particularly useful for functional BGC characterization and prioritizing candidate genes within a cluster of interest [45]. Such multi-omics approaches have been key in recent discovery efforts in biosynthesis [46], and in the development of tools like MEANtools [47], NPLinker [48], and other integrative approaches [49,50].

Understanding the evolutionary significance and biological rationale behind gene clustering in plants remains a critical area of research. The continued development of tools such as plantiSMASH not only aids in the discovery of valuable natural products and their biosynthetic pathways but also contributes to a deeper fundamental understanding of genomic organization and its functional implications in plants. As antiSMASH has become the cornerstone of the ecosystem of computational tools around natural product research, we anticipate that this plantiSMASH update will foster collaborations for the development of compatible tools for plant natural product discovery and biosynthesis prediction.

## Code and Data availability

The plantiSMASH 2.0 web app is available at https://plantismash.bioinformatics.nl/. The plantiSMASH documentation is available at https://plantismash.github.io/documentation. The plantiSMASH database is available at https://plantismash.bioinformatics.nl/precalc/, including TFBS Finder results for *A. thaliana* (https://plantismash.bioinformatics.nl/precalc/v2/tfbs-finder/). These websites are free and open to all users, and there is no login requirement. The plantiSMASH code is open source and is freely available at https://github.com/plantismash/plantismash. All data related to plantiSMASH are deposited in the plantiSMASH Zenodo community (https://zenodo.org/communities/plantismash/), including the clusterBLAST database (https://doi.org/10.5281/zenodo.16927684). The list of curated TFBS motifs, including the data used for their validation, is available at https://doi.org/10.5281/zenodo.17144325. The data for the validation of TFBS detection was originally downloaded from https://connectf.s3.amazonaws.com/connectf_data_release_v1.tar.gz.

## Supplementary Information

The research data and code associated with this publication are available as described in **Code and Data availability** and the **Supplementary Material**.

### Supplementary tables

– **Table S1**: Collection of BGCs curated from literature
– **Table S2**: BGC rules in plantiSMASH 2.0
– **Table S3**: PFAM domains included in plantiSMASH 2.0
– **Table S4**: Genomes included in the database and BGC counts in plantiSMASH version 1.0 and 2.0

## Author contributions

**EDP:** Conceptualization, Data curation, Formal analysis, Methodology, Validation, Visualization, Writing—original draft. **CO:** Conceptualization, Data curation, Formal analysis, Validation, Visualization, Writing—review & editing. **ZL:** Data curation, Formal analysis, Validation, Visualization, Writing—review & editing. **HEA:** Methodology, Validation, Visualization, Writing— review & editing. **AD:** Software, Validation. **GP:** Methodology, Writing—review & editing. **SAK:** Methodology, Software. **AO:** Conceptualization, Methodology. **JJJvdH:** Conceptualization, Project administration, Supervision, Writing—review & editing. **MHM:** Conceptualization, Funding acquisition, Methodology, Project administration, Supervision, Writing—review & editing.

## Acknowledgments

We thank the MIBiG collaborators, including Mitja Zdouc, and the contributors to the plant BGC annotation efforts, in particular Adriano Rutz and Tito Damiani. We also thank our antiSMASH collaborators at the Technical University of Denmark—Kai Blin, Simon Shaw, and Tilmann Weber—for technical support and feedback on the plantiSMASH documentation. We thank Dirk-Jan M. van Workum for assistance with documentation and installation, and Esteban Charria Girón for proofreading.

