## Supplementary Material for "plantiSMASH 2.0: improvements to detection, annotation, and prioritization of plant biosynthetic gene clusters"

**SUPPLEMENTARY INFORMATION**

Elena Del Pup^1^, Charlotte Owen^3†^, Ziqiang Luo^1†^, Hannah E. Augustijn^1,2^, Arjan Draisma^1^, Guy Polturak^4^, Satria A. Kautsar^5^, Anne Osbourn^3^*, Justin J.J. van der Hooft^1^*, Marnix H. Medema^1,2^*

^1^ Bioinformatics Group, Wageningen University, Wageningen, The Netherlands
^2^ Institute of Biology, Leiden University, Leiden, The Netherlands
^3^ John Innes Centre, Colney, Norwich, UK
^4^ The Robert H. Smith Institute of Plant Sciences and Genetics in Agriculture, The Hebrew University of Jerusalem, Herzl 229, Rehovot 7610001, Israel
^5^ DOE Joint Genome Institute, Lawrence Berkeley National Labs, Berkeley, CA 94720, USA

† Joint second authors.

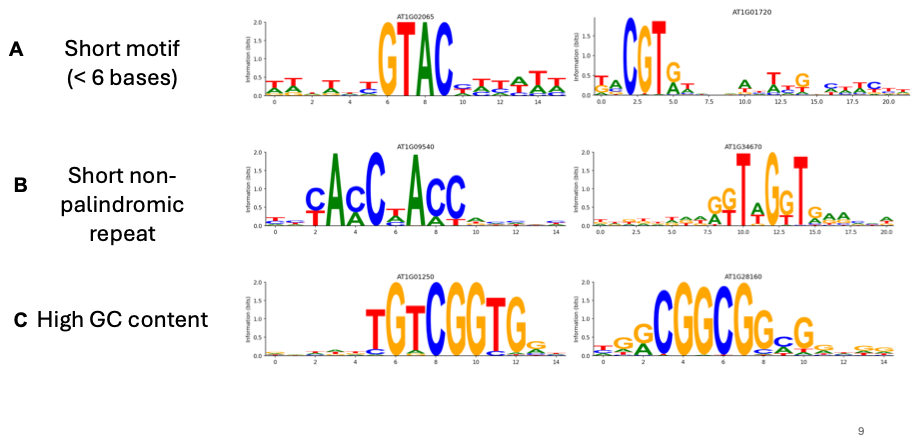

**Fig. S1**

Examples of motifs from the exclusion list of TFBS motifs resulting from manual curation. Motifs are available in PlantTFDB (<https://planttfdb.gao-lab.org/>). The full motif exclusion list is available at <https://doi.org/10.5281/zenodo.17144325>.

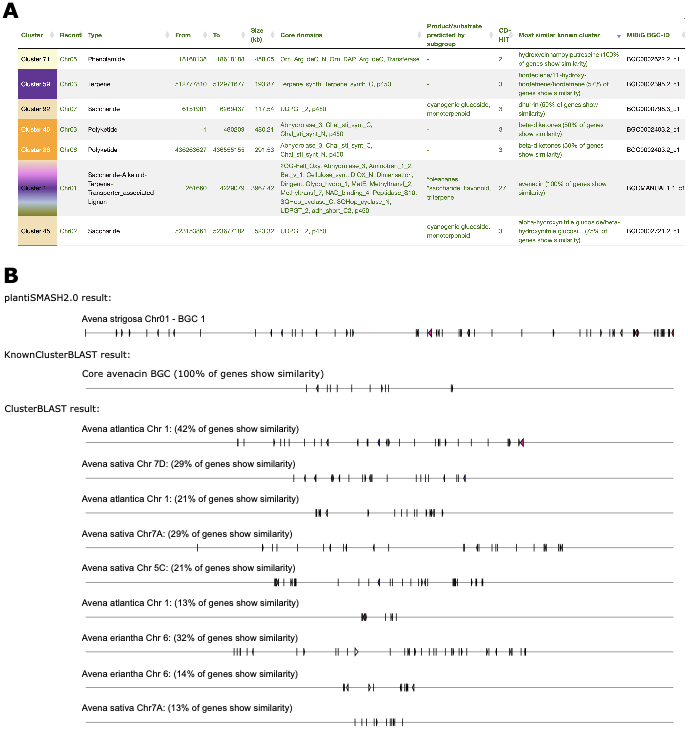

**Fig. S2**

BGCs from *Avena strigosa* with knownclusterBLAST hits. A) knownclusterBLAST shows a putative phenolamide cluster in chromosome 5 homologous to the hydroxycinnamoylputrescine phenolamide gene cluster in *Oryza sativa* Japonica group (MIBiG entry BGC0002622.2). A 75% amino acid similarity was also found for a putative saccharide cluster on chromosome 2 to the α-hydroxynitrile glucoside biosynthetic gene cluster from *Hordeum vulgare subsp. vulgare* (BGC0002721). Partial similarity is also found in the genome of A. strigosa for a putative polyketide and phenolamide BGC with the beta-diketones BGC (BGC0002403) from *H. vulgare subsp. vulgare*, for a putative terpene BGC with hordediene biosynthetic genes (BGC0002395) in *H. vulgare subsp. vulgare*, and for a saccharide BGC against the dhurrin BGC (BGC0000798) in *Sorghum bicolor*. B) ClusterBLAST shows homologous hits of the avenacin BGC to other Avena species, such as *A. eriantha*, *A. atlantica*, and *A. sativa*.

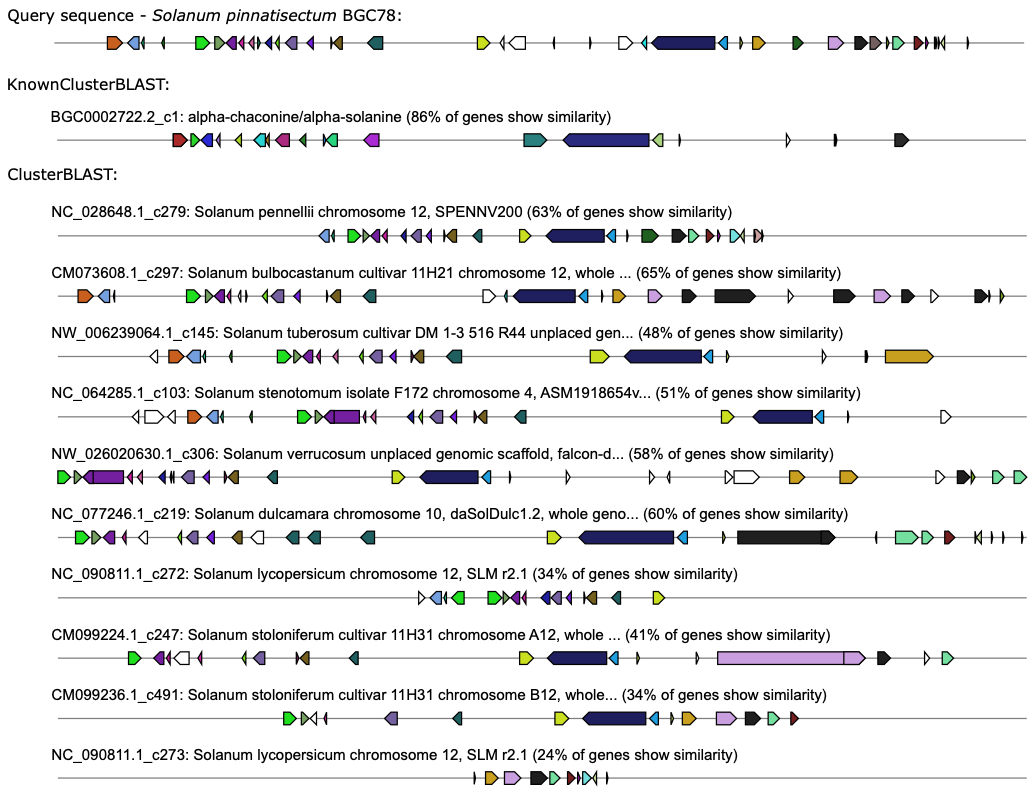

**Fig. S3**

ClusterBLAST output for BGC78 from *Solanum pinnatisectum*.

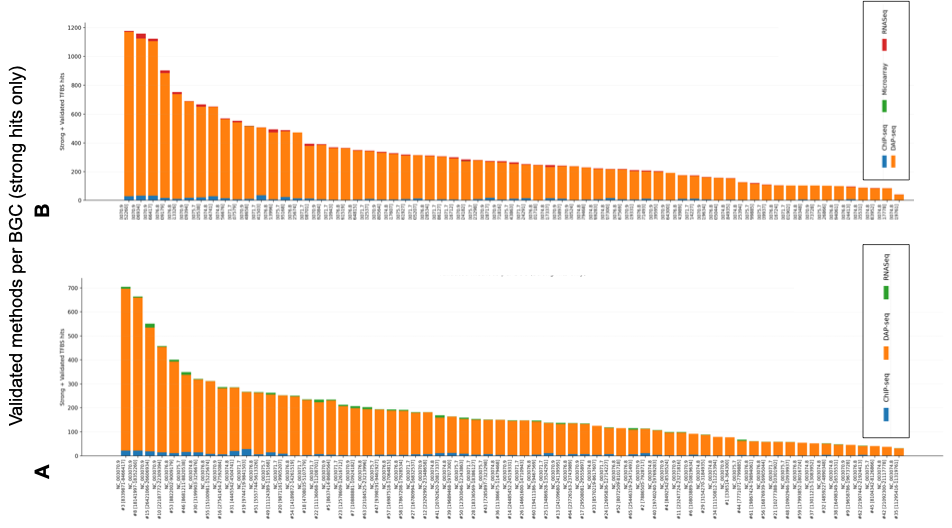

**Fig. S4**

Validated TFBS motifs by Biosynthetic Gene Cluster (BGC) by experimental method in *Arabidopsis thaliana* for strong hits (score ≥ (min_score + max_score)/2). Left: strong hits per validation method across 354 manually curated motifs. Right: strong hits per validation method across 576 motifs.

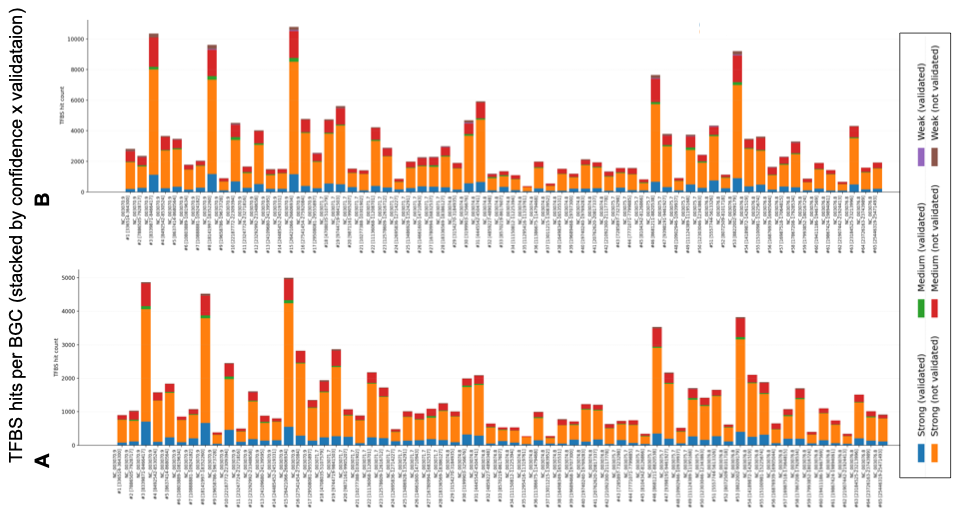

**Fig. S5**

TFBS motifs by Biosynthetic Gene Cluster (BGC) divided by weak (score ≤ min_score), medium and strong (score ≥ (min_score + max_score)/2) hits, including validated ones. Left: 86,111 total hits and 12,475 validated hits across 354 manually curated motifs. Right: 190,676 total hits and 26,276 validated hits across 576 motifs.

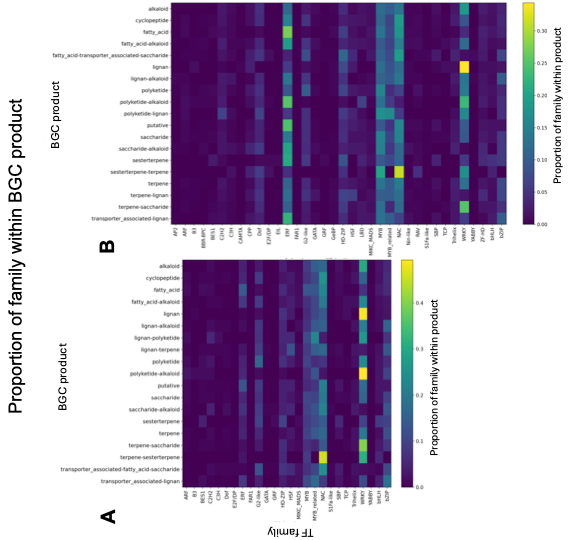

**Fig. S6**

Transcription Factor Binding Site (TFBS) motifs (strong hits validated with ChIP-seq) by cluster product type and TF family from PlantTFDB classification. The heatmap contains the row-normalized TF family proportions per product. Left: enrichment analysis across 354 manually curated motifs. Right: enrichment analysis across 576 motifs.

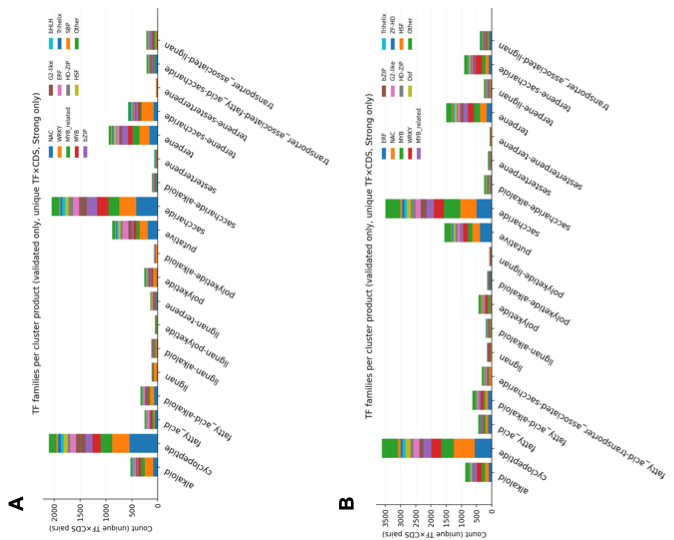

**Fig. S7**

Transcription Factor Binding Site (TFBS) motifs (strong hits validated with ChIP-seq) by TF family for the top 12 TF families across Biosynthetic Gene Cluster (BGC) types. Top: enrichment analysis across 354 manually curated motifs. Bottom: enrichment analysis across 576 motifs.
